# Harnessing ALDH1A2 vulnerability in T-cell acute lymphoblastic leukemia

**DOI:** 10.1101/2025.03.11.642712

**Authors:** Guillaume P. Andrieu, Lucien Courtois, Mickael Bonnet, Camille Gillet, Mélodie Quéri, Mélanie Féroul, Ludovic Lhermitte, Patrick Villarèse, Aurélie Cabanes-Hamy, Phillippe Rousselot, Mileydis Perez, Ismail Ceylan, Salvatore Spicuglia, Vahid Asnafi

**Affiliations:** Institut Necker-Enfants Malades (INEM), INSERM U1151 CNRS UMR8253, Université Paris Cité, Paris, France; Laboratory of Onco-Hematology, Assistance Publique-Hôpitaux de Paris (AP-HP), Hôpital Universitaire Necker Enfants-Malades, Université Paris Cité, Paris, France; Service d’Hématologie et d’Oncologie, Hôpital Universitaire de Versailles, APHP, Versailles France; Advanced BioDesign, Lyon, France; Aix-Marseille University, INSERM, TAGC, UMR 1090, Marseille, France

## Abstract

T-cell acute lymphoblastic leukemia (T-ALL) is an aggressive hematologic malignancy with limited therapeutic options, particularly in the relapsed/refractory (R/R) setting. Unlike other hematological malignancies, which benefit from targeted immunotherapies, T-ALL remains reliant on chemotherapy, leading to poor outcomes in R/R cases. Identifying novel therapeutic vulnerabilities is crucial to improving the outcomes of patients. Herein, we identify aldehyde dehydrogenase 1A2 (ALDH1A2) as a T-ALL-specific enzyme essential for leukemic cell survival. Transcriptomic and epigenetic analyses reveal its selective expression, regulated by the TAL1 oncogene. Pharmacological inhibition of ALDH1A2 using Dimate demonstrates potent anti-leukemic activity across diverse T-ALL subtypes, including primary samples of relapsed disease. These findings establish ALDH1A2 as a therapeutic target in T-ALL and support ALDH inhibition as a promising strategy to overcome drug resistance and improve treatment outcomes in R/R T-ALL.

## Introduction

Acute lymphoblastic leukemia (ALL) is the most frequent leukemia in children, accounting for 30% of the pediatric cancers. These aggressive hematological malignancies also occur in adolescents and young adults. Over the past decades, major improvements were obtained in the treatment of ALL, with the implementation of modern polychemotherapy and revolutionary immunotherapies. Yet, these breakthroughs unevenly benefited the patients suffering from different ALL subtypes. While B-cell ALL patients receive frontline immunotherapies and can have access to anti-CD19 CAR T cells when they develop a refractory or relapsed disease, these major therapeutic options are not available for patients suffering from T-cell ALL (T-ALL). While being actively investigated, immunotherapies are not accessible to T-ALL patients due to safety limitations that represent long-term T-cell aplasia and biological constraints such as the fratricide effect, a major hurdle to T-ALL-reactive CAR T cells. As a result, the therapeutic arsenal of T-ALL mostly relies on polychemotherapy and emerging targeted therapies. In about 30-40% of the cases, these regimens fail to obtain durable complete remission for the patients who develop a refractory disease or experience relapse (R/R T-ALL). The prognosis of these R/R patients is grim, with a 5-year overall survival inferior to 30% (Eckert *et al*., 2021). Hence, it is urgent to address the unmet clinical need of the underserved T-ALL patients by implementing novel targeted therapies with strong anti-leukemic potential against R/R disease.

A better comprehension of the biological liabilities of T-ALL blasts has led to promising targeted therapies entering the clinics (Courtois *et al*., 2023; Pinton *et al*., 2024; Andrieu *et al*., 2025). Notably, the targeting of detoxifying mechanisms involved in chemotherapy derivatives clearance appears as a promising option to eradicate persister cancer cells. Aldehyde dehydrogenases (ALDH) are a family of enzymes involved in cellular detoxification by oxidizing aldehydes into less reactive carboxylic acids. These enzymes play crucial roles in drug metabolism, resistance mechanisms, and stemness properties of cancer cells (Fleischman Blood 2012). Recent studies shed light on ALDH functions in T-ALL, suggesting that these enzymes may constitute a targetable vulnerability of T-ALL blasts (Longville *et al*., 2015; Venton *et al*., 2016; Zhang *et al*., 2021).

## Results

### ALDH profiling confirms ALDH1A2 as a T-ALL isoenzyme

The interrogation of public transcriptomic data collected on pan-cancer cell lines confirmed the elevated expression of ALDH1A2 in T-ALL regarding other hematological malignancies and solid tumors (Figure 1A). Despite the heterogeneous expression across the different cell lines, T-ALL appears at one of the cancers expressing the most ALDH1A2. We further examined a large series of T-ALL primary diagnostic samples profiled by integrated multiomics for the profile of expression of the 19 enzymes of the ALDH subfamily. Transcriptomics revealed ALDH1A2 as a T-ALL-specific isoenzyme, while the other members were not detected across the profiled series. These results are in line with recent studies that have shown that the expression of *ALDH1A2* is driven by the T-ALL major oncogene TAL1 (Longville *et al*., 2015; Zhang *et al*., 2021). Importantly, while TAL1-dependent mechanisms of regulation have been proposed, the interrogation of a large series of T-ALL primary samples revealed that ALDH1A2 is expressed at various levels across other molecular subtypes, in addition to TAL1-positive cases (Figure 1C). To further evaluate the regulation of ALDH1A2 expression in T-ALL, we interrogated a series of 155 T-ALL primary samples profiled by EPIC for DNA methylation. Interestingly, the *ALDH1A2* promoter region is found hypermethylated in T-ALL when compared to normal thymocytes (Figure 1D). Yet, these regions are heterogeneously methylated across the diverse T-ALL subtypes. *ALDH1A2* promoter region methylation follows the global pattern of gene regulatory elements methylation of T-ALL that we unraveled (Figure 1E) (Touzart *et al*., 2021). TAL1-positive T-ALL display low *ALDH1A2* promoter region methylation in line with the high expression of the enzyme detected in these cases (Figure 1F-G). TAL1+ T-ALL is the majority of the C2 cluster previously reported (Touzart *et al*., 2021). Of note, T-ALL with hypomethylation of the regulatory elements of *ALDH1A2* are also found among the TLX1, TLX3 and negative oncotype groups. These cases exhibit high expression of ALDH1A2 (Figure 1C). Collectively, these observations confirm ALDH1A2 as a T-ALL isoenzyme and strongly suggest a functional dependence to its activity.

**Figure 1.**
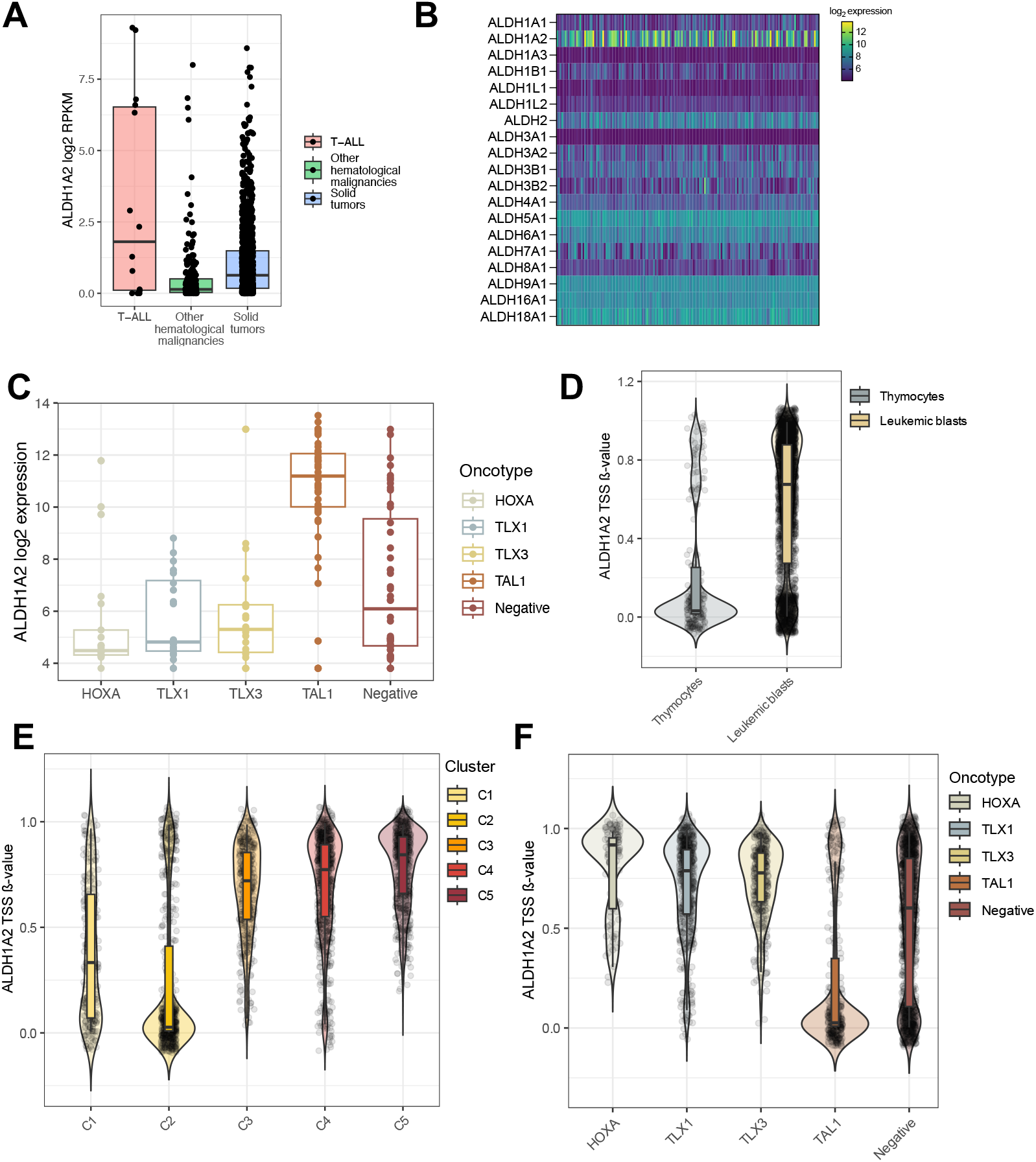
ALDH profiling confirms ALDH1A2 as a T-ALL isoenzyme. A. Exploration of pan-cancer ALDHA12 expression from the DepMap database. B. Profile of expression of ALDH isoenzymes from the Necker T-ALL series (n=155). C. *ALDH1A2* expression stratified by T-ALL oncotypes. D ALDH1A2 TSS DNA methylation in normal thymocytes and T-ALL samples. E-F. ALDH1A2 TSS DNA methylation in T-ALL samples stratified by EPIC clusters (Touzart *et al*., 2021) (E) or by oncotype (F).

### ALDH inhibition exhibits robust anti-leukemia efficacy

To ascertain the reliance of T-ALL blasts on ALDH1A2, we evaluated the efficacy of dimethyl ampal thiolester (Dimate), an ALDH inhibitor with high affinity regarding the ALDH1A2 isoenzyme (Venton *et al*., 2016) in a series of 20 T-ALL patient-derived xenografts of primary samples (PDX) treated *ex vivo*. Strikingly, ALDH1A2 inhibition demonstrated robust anti-leukemic effect on all the tested samples (Figure 2A). Interestingly, 13 out of the 20 samples (65%) presented a strong response at doses inferior to the calculated IC50 for the series (mean IC50: 2.68 µM, median IC50: 1.84 µM, n = 20). A second group of samples (7/20, 35%) presented a response to higher doses, with no clear refractoriness. Dimate efficacy was observed irrespective of the phenotypic or the molecular subgroups of the tested samples (Figure 2C). Interestingly, despite higher expression in TAL1+ T-ALL, no different sensitivity to Dimate was observed among TAL1+ and TAL1-cases (Figure 2 D-E), raising the question of the correlation between the expression and the functional dependency to the enzyme.

**Figure 2.**
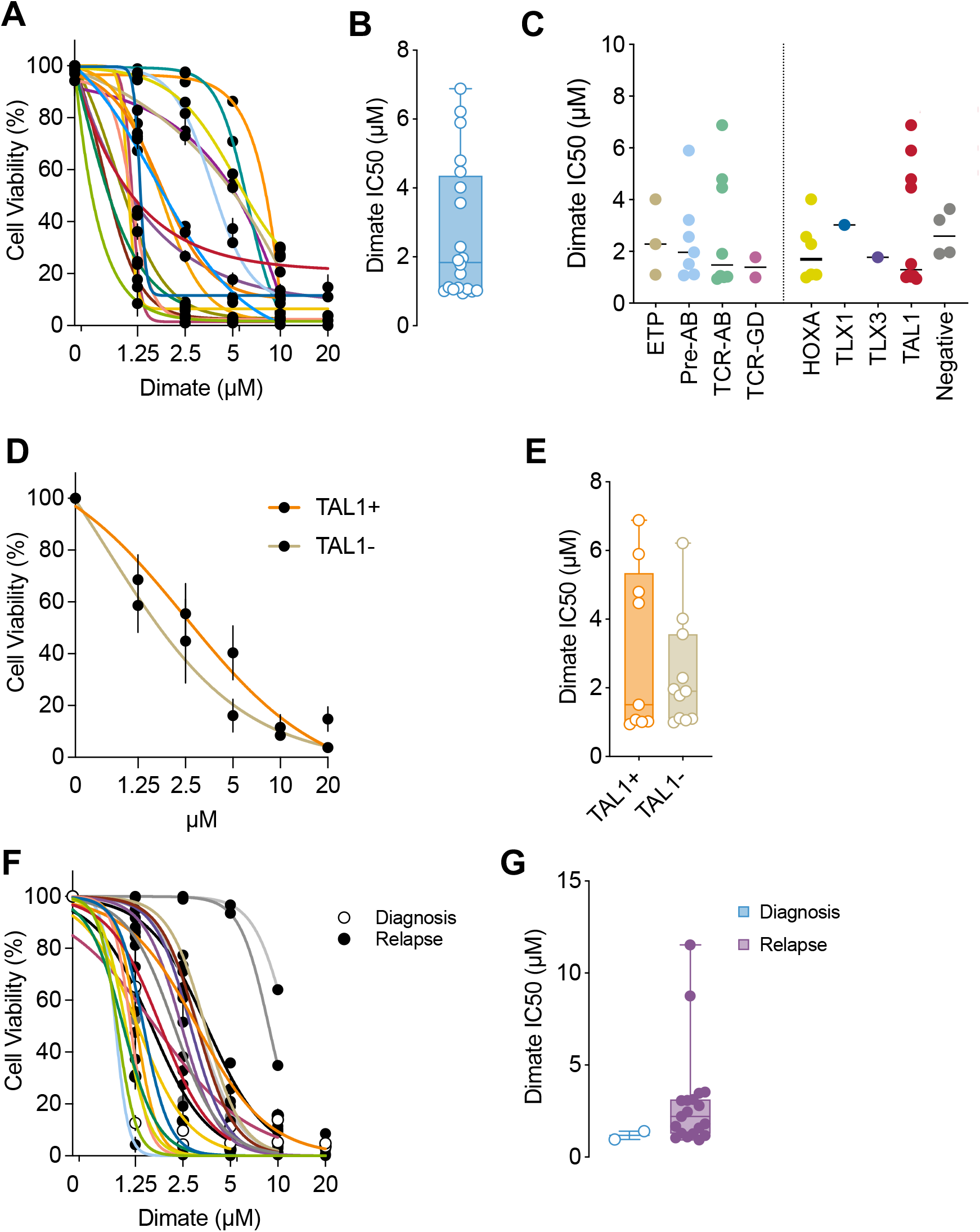
ALDH1A2 inhibition exhibits robust anti-leukemia efficacy. A. Dimate efficacy evaluated *ex vivo* on 20 T-ALL PDX at 72h by flow cytometry-based measure of cell viability. B. IC50 of Dimate for each PDX (n=20). C. Dimate IC50 stratified by phenotype or oncotype. D. Dimate efficacy in TAL1+ and TAL1-PDX (n=20). E. Dimate IC50 in TAL1+ and TAL1-PDX (n=20). F. Dimate efficacy evaluated *ex vivo* on primary samples (n=2 diagnosis, n=20 relapse) at 72h by flow cytometry-based measure of cell viability. G. IC50 of Dimate for each primary sample (n=2 diagnosis, n=20 relapse).

To further explore the anti-leukemic effect of ALDH1A2 inhibition, we treated a series of 20 T-ALL primary samples collected at relapse. Strikingly, Dimate demonstrated robust cytotoxicity on these refractory samples (IC50: 2.78 µM, Figure 2F-G). Similar effects were obtained on 2 primary samples obtained at diagnosis. Of note, two relapses with an ETP phenotype with *ABL* rearrangements were found less sensitive to the compound. Altogether, these data obtained on primary samples and derived xenografts illustrate the potent anti-leukemic activity of Dimate, suggesting that ALDH1A2 inhibition may represent an efficient strategy to treat T-ALL, including in relapsed/refractory settings.

To validate this, we tested Dimate *in vivo* on an aggressive TAL1+ T-ALL PDX that presented sensitivity *ex vivo* (IC50: 1.07 µM). Because of its structural properties, Dimate requires lipid encapsulation to prevent its rapid degradation *in vivo*. Once the engraftment of the PDX confirmed by a blood infiltration of 0.5% to 5%, mice were treated either with vehicule (ABD-3000, lipid carrier only) or encapsulated Dimate (ABD-3001, 5 mg/kg) by intraperitoneal injections three times a week for 24 days (Figure 3A). We noted no toxicity for the animals which maintained stable weight on treatment (Figure 3B). While leukemia aggressively propagated in the vehicule group, Dimate controlled leukemia outgrowth over the course of treatment, except for one animal (Figure 3C). The monitoring of the leukemic burden in these animals revealed the expansion of the disease after stopping the treatment. As a result, Dimate-treated animals presented a promising extension of survival (Figure 3D).

**Figure 3.**
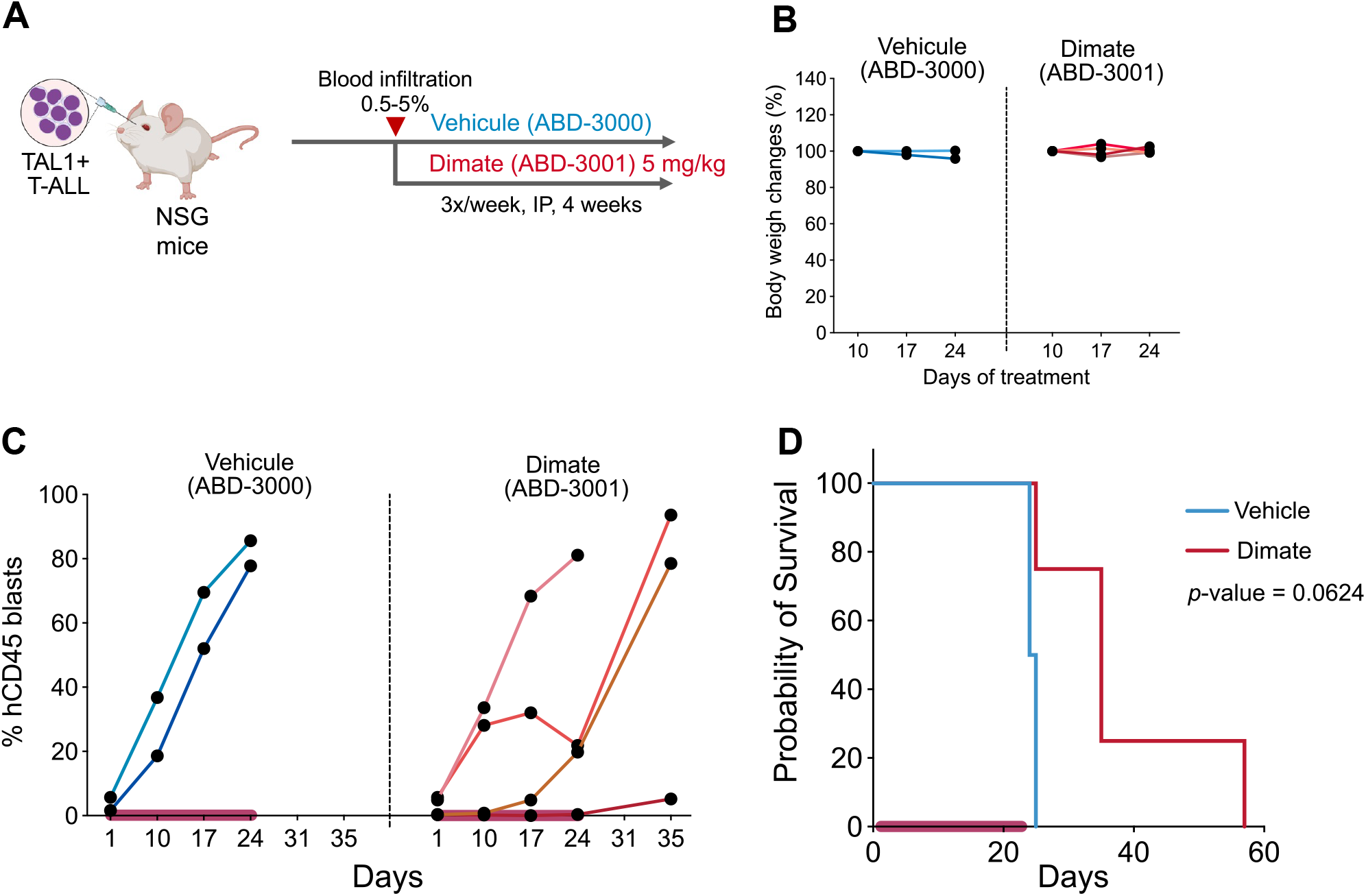
Efficacy of encapsulated Dimate *in vivo*. A. Design of the experiment. B. Animal weight follow-up on therapy. C. Tumor burden in vehicule and Dimate-treated animals. The pink ribbons indicate the duration of treatment. D. Animal survival curves of vehicule and Dimate groups. The exact p-value is indicated. The pink ribbon indicates the duration of treatment.

Collectively, these results shed light on the T-ALL dependency to ALDH1A2. We provide *ex vivo* an *in vivo* data sustaining ALDH1A2 inhibition as a promising targeted therapy to counter T-ALL. Critically, we report unprecedented single-agent efficacy against primary samples of relapsed disease that failed standard-of-care therapy, underpinning the interest of ALDH inhibitors in the treatment of lethal R/R T-ALL.

## Discussion

The clinical challenge constituted by R/R T-ALL patients is worrisome. The limited therapeutic options available for these patients is a critical hurdle to overcome. As in 2025, nelarabine is the only approved molecule to treat R/R T-ALL, with limited efficacy (Gökbuget *et al*., 2012). Hence, the unmet clinical needs of R/R T-ALL underscore the urgency of identifying and validating novel targeted therapies that exploit the specific vulnerabilities of leukemic blasts. Notably, the inhibition of the cellular mechanisms upregulated in chemopersister cells may constitute promising options.

The overexpression of ALDH enzymes is a hallmark of cancer (Dinavahi *et al*., 2019). Several studies reported that chemotherapy- and radiotherapy-resistant cancer cells overexpress various members of the ALDH family. Aldehydes are derivatives of the metabolism of amino acids, lipids, vitamins, or alcohols, and are also produced from oxidative stress and chemotherapy (Rodríguez-Zavala *et al*., 2019). Hence, ALDH are essential actors of the anti-oxidant responses that cancer cells engage to resist stressful environmental conditions. In solid tumors, the roles of diverse ALDH have been recently described. The expression of ALDH1 isoenzymes conveys a prognosis value in skin, lung, breast, prostate, colorectal and digestive cancers (Dinavahi *et al*., 2019). In addition, ALDH1 plays critical functions in stem cell biology, notably *via* the production of retinoic acid (Ma and Allan, 2011). ALDH1 has been proposed as a marker for cancer stem cells in solid and hematological tumors (Ginestier *et al*., 2007; Lavudi *et al*., 2024). Several studies have coined the role of ALDH enzymes in chemoresistance (Januchowski, Wojtowicz and Zabel, 2013; Lavudi *et al*., 2024).

We confirm a growing body of evidence identifying ALDH1A2 as a T-ALL isoenzyme in a large series of primary samples. The ALDH family comprises 19 members, with some overlapping substrates and functions. In T-ALL, ALDH1A2 appears to be the sole ALDH expressed, suggesting a strong dependency on this isoenzyme (Longville *et al*., 2015; Zhang *et al*., 2021). In line with this, ALDH1A2 inhibition presented a robust anti-leukemic effect, even on refractory or relapsed blasts. ALDH1A2 expression is regulated by TAL1, a major driver oncogene of T-ALL. As a result, TAL1+ T-ALL is the subgroup with the highest ALDH1A2 expression. Strikingly, ALDH1A2 inhibition was efficient across all the subgroups of the disease, and TAL1+ T-ALL did not exhibit higher sensitivity. Hence, ALDH1A2 expression may not reflect the response to Dimate, which raises questions on its mechanisms of action and biomarkers of response.

The important biological functions exerted by ALDH in cancer stem cells and persister cells combined with the unique pattern of expression of ALDH1A2 in T-ALL strongly suggests that Dimate may have potent cytotoxicity against these chemorefractory blasts that resist therapy and drive lethal disease. Further evaluation is required to position ALDH1A2 inhibition as an efficient strategy to target leukemia chemopersister cells and prevent relapse.

## Methods

### Patient-derived xenografts

NOD.Cg-*Prkdc*^SCID^ *Il2rg*^tm1Wjl^/SzJ (NSG) mice were purchased from Charles River and maintained in a specific and opportunistic pathogen-free LEAT Animal facility at the Institut Necker-Enfants Malades. Patient-derived xenografts (PDX) were generated as previously described (Andrieu *et al*., 2021, 2025; Touzart *et al*., 2021). Briefly, 10^6^ fresh viable primary blasts were xenografted by intravenous retro-orbital injection in 6-week-old NSG mice. The leukemic burden in the blood was monitored weekly by flow cytometry as the percentage of FSC^hi^, hCD45^+^ cells. Mice were euthanized when terminally ill, as evidenced by a blood infiltration >80%, severe dyspnea or frailty caused by leukemic dissemination in the mediastinum or vital organs, by the IRB approval. The maximal tumor burden of 90% blood infiltration was not exceeded. For *ex vivo* experimentation, bone marrows from tibiae, hips, femurs, and vertebrae were collected and blasts purified by Ficoll. The blast purity was checked by flow cytometry (percentage of FSC^hi^, hCD7^+^ hCD45^+^ cells) before experimentation. All the samples used contained ≥90% human blasts.

For *in vivo* experimentation, mice were segregated into treatment arms when the leukemic load reached 0.5% to 5% (5 mice per arm) and treated for vehicule (ABD-3000) or encapsulated Dimate (ABD-3001, 5 mg/kg) by intraperitoneal injections three times a week for 24 days.

### *Ex vivo* ALDH1A2 inhibition

*Ex vivo* cultures of PDX were achieved in a complete medium, supplemented with 50 ng/ml human stem cell factor, 20 ng/ml hFLT3-L, 10 ng/ml hIL-7, and 20 nM insulin (Miltenyi Biotec, Bergisch Gladbach, Germany) and 20% fetal bovine serum. Cultures were maintained at 37°C in a humidified atmosphere containing 5% CO_2_. Dimate and its encapsulated *in vivo* formulation (ABD-3001), and the vehicule (ABD-3000) were obtained from Advanced BioDesign.

### RNA sequencing and data analysis

155 primary samples were analyzed by RNA sequencing. RNA eligibility was determined by measurement on a BioAnalyzer (RIN ≥ 8, Agilent). Libraries were prepared using the SureSelect XT HS2 kit (Agilent) and sequenced on a NovaSeq sequencer (Illumina). Reads were trimmed using Agent v2.0.5 with ‘trim -v2’ arguments and aligned to the GRCh38 genome assembly using STAR v2.7.9 with default parameters. before deduplication using Agent with ‘locatit mbc -i -R’ arguments. Gene-level counting was done using subread featureCounts v2.0.0 with -O argument against Ensembl release 95 gtf file.

Public pan-cancer data on ALDH1A2 expression were retrieved from DepMap.

### DNA methylation analysis

DNA methylation using Infinium Methylation EPIC BeadChip (Illumina, San Diego, CA, USA) on primary T-ALL samples was performed following the manufacturer’s protocol as previously reported (Touzart *et al*., 2021).

### Statistical analyses

Normality tests were applied to determine if the datasets were eligible for either parametric or non-parametric tests. Statistical analyses were performed with Student *t*-test, ANOVA, or Fisher test according to the dataset nature by using GraphPad Prism 8 software (GraphPad Software, Inc., San Diego, CA, United States) and R (v. 4.3.1). IC50 values were calculated by performing non-linear regression on the raw data with GraphPad Prism 8. Animal survival curves were analyzed by log rank Mantel-Cox test.

## References

Andrieu, G.P. et al. (2021) ‘PRC2 loss of function confers a targetable vulnerability to BET proteins in T-ALL’, Blood, 138(19), pp. 1855–1869. Available at: 10.1182/blood.2020010081.

Andrieu, G.P. et al. (2025) ‘A metabolic synthetic lethality of phosphoinositide 3-kinase-driven cancer’, Nature Communications, 16(1), pp. 1–15. Available at: 10.1038/s41467-025-57225-7.

Courtois, L. et al. (2023) ‘IL7-receptor expression is frequent in T-cell acute lymphoblastic leukemia and predicts sensitivity to JAK-inhibition’, Blood, p. blood.2022017948. Available at: 10.1182/blood.2022017948.

Dinavahi, S.S. et al. (2019) ‘Aldehyde Dehydrogenase Inhibitors for Cancer Therapeutics’, Trends in Pharmacological Sciences, 40(10), pp. 774–789. Available at: 10.1016/j.tips.2019.08.002.

Eckert, C. et al. (2021) ‘Risk factors and outcomes in children with high-risk B-cell precursor and T-cell relapsed acute lymphoblastic leukaemia: combined analysis of ALLR3 and ALL-REZ BFM 2002 clinical trials’, European Journal of Cancer, 151, pp. 175–189. Available at: 10.1016/j.ejca.2021.03.034.

Ginestier, C. et al. (2007) ‘ALDH1 Is a Marker of Normal and Malignant Human Mammary Stem Cells and a Predictor of Poor Clinical Outcome’, Cell Stem Cell, 1(5), pp. 555–567. Available at: 10.1016/j.stem.2007.08.014.

Gökbuget, N. et al. (2012) ‘Outcome of relapsed adult lymphoblastic leukemia depends on response to salvage chemotherapy, prognostic factors, and performance of stem cell transplantation’, Blood. 2012/04/12 edn, 120(10), pp. 2032–2041. Available at: 10.1182/blood-2011-12-399287.

Januchowski, R., Wojtowicz, K. and Zabel, M. (2013) ‘The role of aldehyde dehydrogenase (ALDH) in cancer drug resistance’, Biomedicine & Pharmacotherapy, 67(7), pp. 669–680. Available at: 10.1016/j.biopha.2013.04.005.

Lavudi, K. et al. (2024) ‘ALDH and cancer stem cells: Pathways, challenges, and future directions in targeted therapy’, Life Sciences, 356, p. 123033. Available at: 10.1016/j.lfs.2024.123033.

Longville, B.A.C. et al. (2015) ‘Aberrant expression of aldehyde dehydrogenase 1A (ALDH1A) subfamily genes in acute lymphoblastic leukaemia is a common feature of T-lineage tumours’, British Journal of Haematology, 168(2), pp. 246–257. Available at: 10.1111/bjh.13120.

Ma, I. and Allan, A.L. (2011) ‘The Role of Human Aldehyde Dehydrogenase in Normal and Cancer Stem Cells’, Stem Cell Reviews and Reports, 7(2), pp. 292–306. Available at: 10.1007/s12015-010-9208-4.

Pinton, A. et al. (2024) ‘PHF6-altered T-ALL Harbor Epigenetic Repressive Switch at Bivalent Promoters and Respond to 5-Azacitidine and Venetoclax’, Clinical Cancer Research, 30(1), pp. 94–105. Available at: 10.1158/1078-0432.CCR-23-2159.

Rodríguez-Zavala, J.S. et al. (2019) ‘Role of Aldehyde Dehydrogenases in Physiopathological Processes’, Chemical Research in Toxicology, 32(3), pp. 405–420. Available at: 10.1021/acs.chemrestox.8b00256.

Touzart, A. et al. (2021) ‘Epigenetic analysis of patients with T-ALL identifies poor outcomes and a hypomethylating agent-responsive subgroup.’, Science translational medicine, 13(595). Available at: 10.1126/scitranslmed.abc4834.

Venton, G. et al. (2016) ‘Aldehyde dehydrogenases inhibition eradicates leukemia stem cells while sparing normal progenitors’, Blood Cancer Journal, 6(9), p. e469. Available at: 10.1038/bcj.2016.78.

Zhang, C. et al. (2021) ‘Oncorequisite role of an aldehyde dehydrogenase in the pathogenesis of T-cell acute lymphoblastic leukemia’, Haematologica, 106(6), pp. 1545–1558. Available at: 10.3324/haematol.2019.245639.

